# Dorsal Premotor Cortex Involved in Hand Gesture Preparation in Macaques

**DOI:** 10.1101/869354

**Authors:** Guanghao Sun, Shaomin Zhang, Ruixue Wang, Yaoyao Hao, Weidong Chen, Xiaoxiang Zheng

## Abstract

Reaching to grasp movement is thought to rely upon two independent brain pathways. The dorsomedial one is involved in reaching while the dorsolateral one is dealing with grasping. However, some recent evidences suggested that the dorsomedial pathway might participate in grasp movement. Therefore, it is important to investigate whether PMd is involved in grasp planning, and if participating, what kind of role PMd played in grasp planning. In this study, two macaques monkeys were trained to grasp same object by instructing or freely choosing one of two grips, power grip or hook grip. A 96-channel microelectrode array was implanted to collect the population activity of PMd in each subject. Both single unit activity and population activity were analyzed. We found that nearly 21.0% and 26.8% units in PMd of two monkeys displayed grip selectivity during gesture planning in both instructing or freely choosing conditions. These units exhibit selectivity for different gestures when facing the identical visual stimuli (freely choosing condition). At the same time, similar activity patterns are displayed for the same gesture when faced with different selection strategies (freely choosing condition vs. instructing condition). These results show that some neurons of PMd are mainly involved in the hand shape preparation and have no obvious relationship with external visual stimuli and selection strategies.

## INTRODUCTION

Reaching to grasp different objects is one of the most commonly used motor functions in activities of daily life (Grafton et al, 2010). This common behavior requires a lot of complex brain operations, such as transporting the hand towards the object and preshaping the hand to match the different size and orientation of the objects. The two-system theory shows that the reach-to-grasp action could be divided into “transport” which is mainly accomplished by the proximal musculature of the forelimb, and “grasp” which is mainly accomplished by the distal musculature (Brochier et al, 2007; Gardner et al., 2007). The two components are controlled by two specific parieto-frontal pathways separately: a dorsomedial pathway which involves the areas of the superior parietal lobule and the dorsal premotor area (PMd), dealing with reaching (Fink et al., 1997; Brrochier et al., 2007; Wise et al., 1997) and a dorsolateral pathway which includes the inferior parietal lobule and the ventral premotor cortex (PMv), involving in grasping (Jeannerod et al., 1995; Jeannerod, 1997; Luppino et al., 1999; Lehmann et al., 2013).

PMd is an important area in the dorsomedial pathway and this area has been extensively studied during the past two decades. Many studies have illustrated that neurons in PMd are correlated to parameters of reaching movements in planning and execution period (Kalaska et al. 1997; Wise et al. 1997). However, these studies have been focused on the contribution of the proximal forelimb movement to the neuronal discharge, the distal forelimb movements were not taken into account. In recent years, several researchers have debated this theory. Raos et al showed that some neurons in PMd were selective for different prehension and orientation in planning and execution period (Raos et al, 2004). Some later studies demonstrated that hand shape, hand dimensions or grasp force of grasp movement could be decoded from the population discharge in PMd which was recorded by intercortical microelectrode array (Stark et al, 2007, 2008; Hendrix et al., 2009; Van et al., 2012; Takahashi, 2017). Hao et al successfully decoded the grasping movement from both spikes and local field potentials of PMd for real-time prosthetic hand control (Hao et al., 2014). In our previous work, we decoded different gestures from PMd in planning period (Sun et al, 2015).

The neuron selectivity in planning period could be caused by external object features or internal hand shape generated. There are some reports showing that PMd has been viewed as a critical node of a parieto-frontal network for visual guidance of reaching movements. Johnson et al and Wise et al. shows that the PMd interconnected with the medial intraparietal area (MIP) which involved in the control of arm reaching movements (Cui & Andersen 2007, Scherberger & Andersen 2007), represents target locations with respect to the direction of gaze and the position of the arm (Johnson et al. 1996, Wise et al. 1997). Cisek & Kalaska find that population activity in PMd responds to a learned visual cue within 50 ms of its appearance in a reaching task (Cisek & Kalaska 2005). Budisavljevic et al demonstrate that the U-shaped premotor connections were significantly related to the visuomotor processing in reaching and reach-to-grasp movements (Budisavljevic et al., 2017).

In contrast, there is powerful evidence that the PMd involves the guidance of internally generated movements. Kurata and Wise find that set-related premotor cortex activity reflects aspects of reaching preparation (Kurata and Wise, 1988). Mitz et al. illustrate that substantial population of premotor cortex neurons showed the predicted learning- dependent changes in activity (Mitz et al., 1991). Ohbayashi et al report that inactivation of the PMd had a marked effect only on the performance of sequential movements that were guided by memory (Ohbayashi et al, 2016). Li et al shows that premotor cortex hemispheres can maintain preparatory activity independently (Li et al., 2016). Ma et al demonstrate that PMd adapt to the external perturbation might reflect a preparation for the impending response (Ma et al., 2016). Kaufman et al find that the neural activity was indecision and vacillation in free choice condition sometimes (Kaufman et al., 2015).

Therefore, in this paper we aimed to distinguish whether the selectivity of neural signals in PMd reflects the behavioral meaning of the cue or encodes the true gesture planning. In addition, if PMd is involved in the gesture planning, its function is biased towards gesture movements or decision-making strategies. Here we designed a free chosen paradigm in which monkeys autonomously chose to grasp one object by either Power or Hook gesture with or without an instruction of right gesture. In this task, we find that part of recorded units in PMd were shown movement selectivity that generalize across the planning condition. Moreover, these units are very similar in terms of quantity and firing pattern under the condition of instruction and no instruction. These results indicate that the PMd is related to internally plan selection and its function is biased towards prepare gesture movements.

## MATERIALS AND METHODS

The care of the monkeys and the experimental protocols complied with the Guide for The Care and Use of Laboratory Animals (China Ministry of Health) and were approved by the Animal Care Committee at Zhejiang University, China.

### A. Behavioral Task

In this study, two adult male rhesus macaques, Monkey B06 and Monkey B09 were trained to grasp one object with two different gestures. Monkeys were seated in front of a transparent plexiglass board, and a handle-shaped object was fixed on the vertical board in front of the monkeys’ chest level. The size of the object was 20 cm × 15 cm × 2.5 cm. There were three touch sensors in the front, side and back of the object, which could identify the grasp types of the monkeys. A button was placed below the object. Three light-emitting diodes (LED), arranged in a downward triangle array, were mounted above the object (Figure 1A). The lower one first signaled Fixation cue and then Go cue when the color changed from red to green. The upper two indicated the Gesture Cue, with red for Power gesture and green for Hook gesture. Figure 1A showed the flowchart of the behavioral tasks. The trial began when the monkey pressed the button. After 300 ms, the Go Signal LED was turned red, and the monkey was required to continue fixating and keep the button pressed until the Go Signal LED turned green. After a delay of 800 ms period, two Gesture Cue LEDs were turned on as three types (showed in figure 1B). In 25% of trials, only the Red Gesture Cue turned on, instructing a Power gesture; the light and gesture were shown at the top of figure 1B. In another 25% of trials, only the Green Gesture Cue turned on, instructing a Hook gesture; the gesture was shown at the bottom of figure 1B. These trials were defined as Instructed condition. In the remaining 50% of trials, both the Red and Green Gesture Cue turned on together. In these trials, the monkey could choose either Power or Hook gesture to grasp the object. These trials were defined as Free-Chosen condition. The Gesture Cue was prompted for a period of 700ms before it turned off. After a Planning period of 600-1000 ms, the color of the Go Signal LED changed to green. After that, the monkey was required to release the button and executed a reach-to-grasp task. The touch sensors in the object verified if the gesture matched the instruction of Gesture Cue. The monkey must hold the object for another 500 ms before Go cue turned off. After that, the monkey was allowed to release the object and prepare for a new trial start. At the end of each correct trial, the monkey was rewarded with some water.

**Figure 1.**
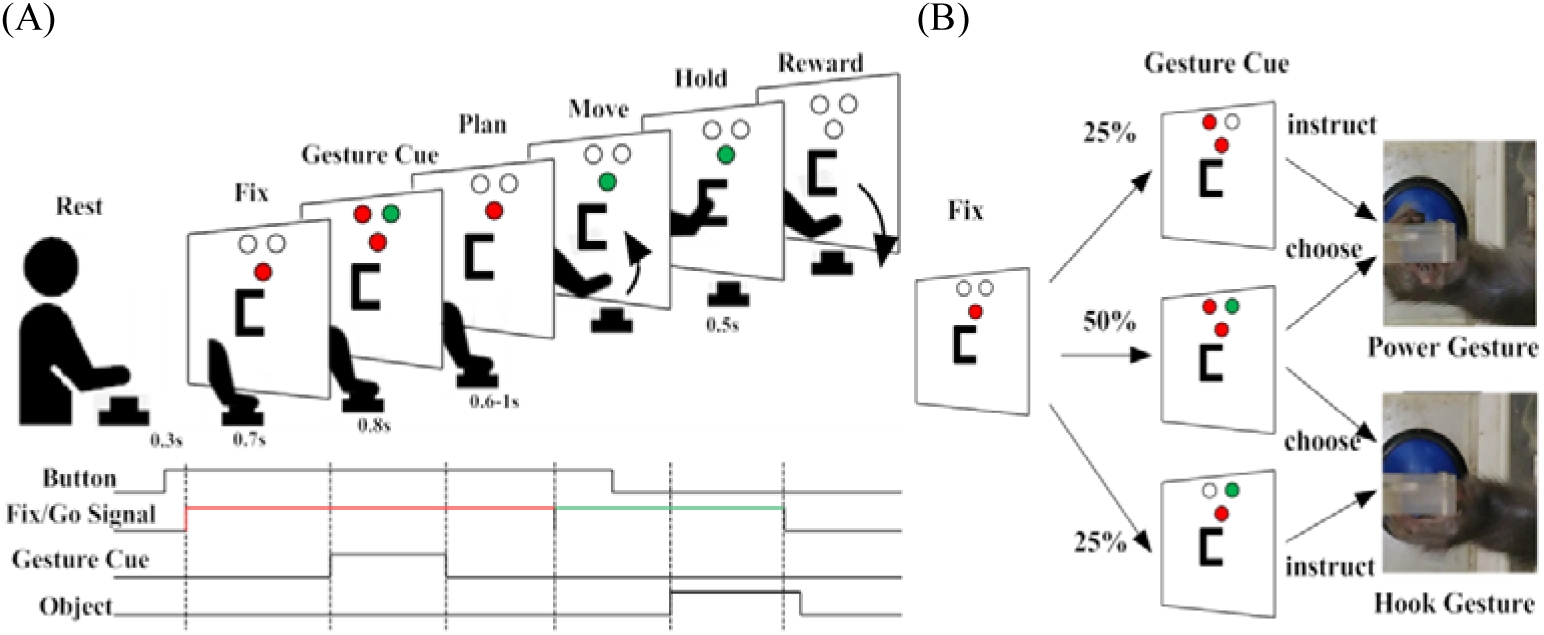
Experimental flow chat and grip types. (A) flow chat and Time course of a trial. The black line upward shows the time that the button has been pressed, gesture cue on and the object has been pressed, the red line marked the LED turn red, and the green line marked the LED turn green; (B) Diagram of Gesture Cue-instructed Power (top) and Hook (bottom) and Free-Chosen trials (middle).

In Free-Chosen trials, the gesture cue information presented to the subject was always the same, but the monkey could freely choose either power or hook gesture to grasp the object. In this way, the selective neurons in PMd (if any) were not caused by external visual information but internal hand shape planning. To make sure that internal hand shape planning was executed after Gesture Cue on, the Free-Chosen trials were interleaved with Instructed trials. Because in Free-Chosen conditions, the monkeys did not know which gesture would be presented before Cue period. The whole task was controlled by a custom program in LabVIEW, which communicated with a DAQ card (NI USB 6001).

### B. Surgery and Data Acquisition

After sufficient training (performance > 85%; for 5-7 months), monkeys randomly chose gesture between Power and Hook which equally happened in the free-chosen trials. Then a microelectrode array (96-channels, 4.0 × 4.0 mm, Blackrock Microsystems, USA) was implanted in PMd area. Figure 2 showed the location of implanted microelectrode array of Monkey B06 and B09. Cerebus multichannel data acquisition system (Blackrock Inc., USA) was used to record the neural signals from PMd. We used the digital input ports of Cerebus to make 9 task events synchronized with the neural signal. The 9 synchronized task events were: Button press, Go Signal Red/Green LED, Gesture Cue Red/Green LEDs, Front/Back/Side Touch Sensors, and Reward. The whole process of the experiments were videotaped with a camera.

**Figure 2.**
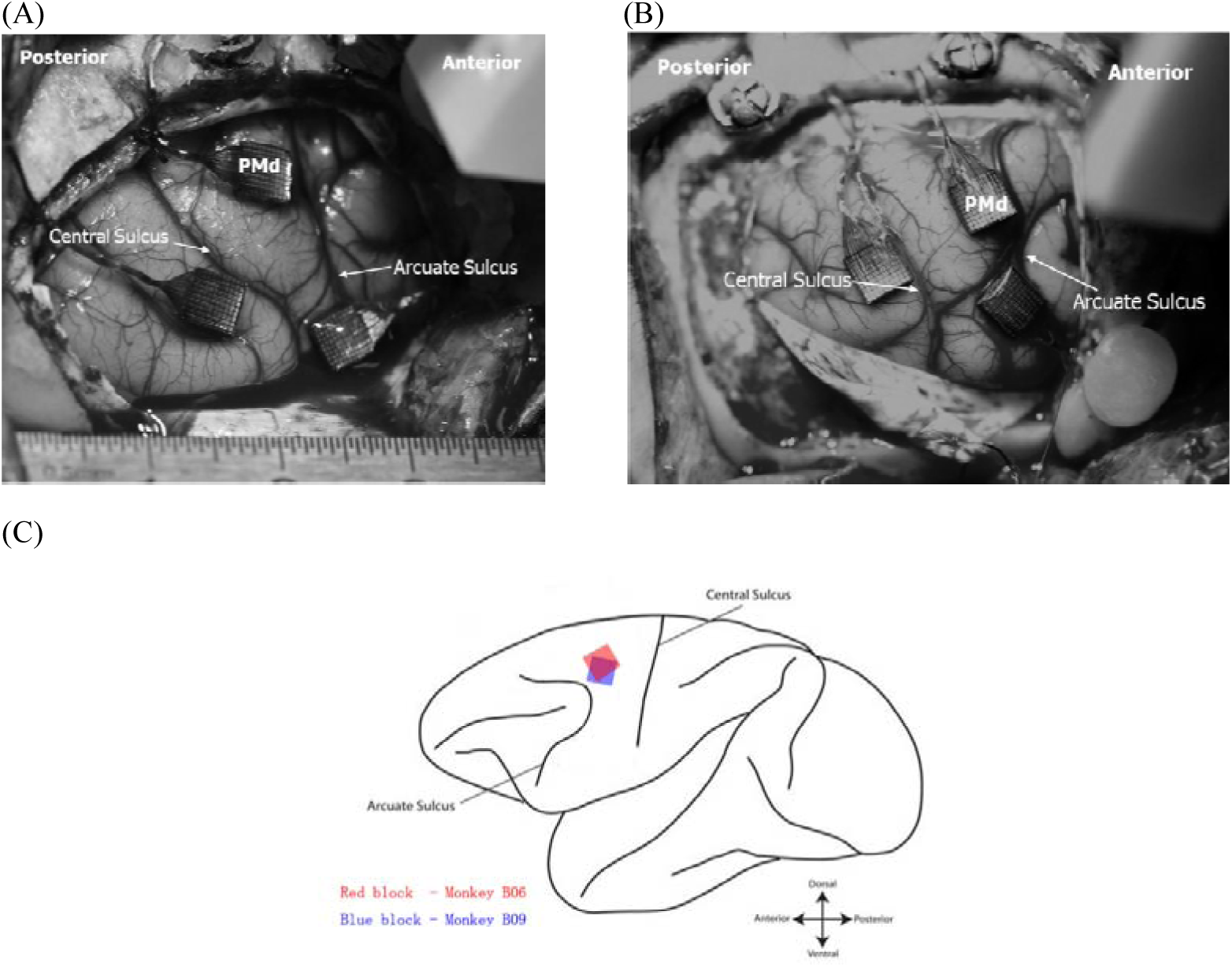
Anterior direction, poster direction, central sulcus and arcuate sulcus are depicted in the photo. (A) Monkey B06. (B) Monkey B09. In this paper, we just analysis the data from PMd area. (C) Sketch map of electrode position

### C. Data Analysis

#### Behavioral data analysis

In Free-Chosen trials, monkeys chose either power or hook gestures to grasp the object autonomously. Both decisions could receive rewards. We should detect whether the monkey showed stereotyped selections or excessively frequent movement choices. First, we calculated a cumulative number of trials in which the monkeys chose Power and Hook gestures in Free-Chosen trials. Then we used Shannon’s Entropy Index to calculate a measure of randomness for gesture choice. The Shannon’s Entropy Index (Shannon, 1948) as follows:

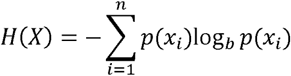

Where *x*_*i*_ is from a transition matrix that shows the number of movement followed any other, and *p*(*x*_*i*_) is the probability mass function of each state *x*_*i*_, and *n* is the number of transition matrix’s states. The range of Shannon’s Entropy Index *H*(*X*) is from 0 to log_2_*n*. A high entropy index means a close probability of each state in transition matrix, and random choice of the monkey between the Power and Hook gestures. In this task, we analyze whether the 1 to 4 history trials affect the current selection.

#### Preprocessing

The spike signals were first extracted from the raw neural data collected by Cerebus system. First, we filtered out the spike waveforms by threshold detection method; the threshold was set at −5.5 times the root mean square (RMS) of baseline signal level. Second, the spike waveforms were projected into the principal component basis to arrive at two coefficients. These coefficients were subjected to manual spike-sorting using Offline Sorter (Plexon Inc., USA). Furthermore, to analyze the tuning characteristics of each unit, we binned the spikes of each unit in non-overlap continuous 100ms windows. Then we extracted the timing of trial events and marked it into the bin data.

#### Data analysis

The analysis of neuronal activity during the Free-Chosen conditions was made by subdividing the discharge into the following epochs: 1) Rest: from 300 ms before Button press to Button press. 2) Press: from Button press to red LED on (300 ms). 3) Fix: from red LED on to Gesture Cue on (700 ms). 4) Cue: from Gesture Cue on to Gesture Cue off. 5) Plan: from Gesture Cue off to Go Signal on. 6) Move: from Button release to the beginning of Object touching.

A neuron was associated with grasp plan when the discharge of neural activity displayed a significant difference between different choice from Cue and Plan period (one-way ANOVA, p<0.01). Peristimulus-time histograms (PSTH) were constructed with power and hook gestures of Free-Chosen condition respectively. The PSTH were smoothed using a Gaussian kernel (half-width of 50 ms). After that we counted the number of units that showed selectivity in both conditions.

Then we calculated the proportion of units that showed selectivity in the Cue, Plan, and Move period in Free-Chosen and Instructed conditions, as well as the proportion of units which showed selectivity in both conditions.

To determine whether neurons exhibited similar responses for forced and free choices, we performed a correlation analysis. For each neuron, we first collected a vector of trial-averaged firing rates over time for the Instructed Power/Hook condition. Firing rates were smoothed with a Gaussian (50 ms SD). We then correlated it (Pearson correlation) with the analogous response vector for the Free-Chosen Power/Hook conditions. The resulting correlation coefficients were averaged over units.

After that, we further investigated the population features of gesture planning in a sliding window analysis. We performed a support vector machines (SVM) to predict results in a 300-ms-width window shifted by 100 ms. In plots, the predicted result of all units and grasp plan related units in each bin was reported at the end position of each window. A 10-folds cross-validation had been used.

The SVM model not only used to predict impending gestures, it will also be used to study the situation of monkeys changing his mind in a single trial. We refer to the work of Kaufman et al., training the SVM using only Instructed trials with delay durations of at least 300 ms. We used only these relatively-long-delay trials so that early delay activity was not overrepresented. After that, the SVM model was used to calculate the gesture choice probabilities of each trial at different times in Free-Chosen and Instructed conditions.

Finally, the Laplacian Eigenmaps (LE) analysis was used to visualize gesture-related effects on neural activity at each period of the task (Belkin et al., 2003). The LE algorithm begins by representation for neural data lying on a low-dimensional manifold embedded in a high-dimensional space. The algorithm provides a computationally efficient approach to nonlinear dimensionality reduction that has locality-preserving properties and a natural connection to clustering.

## RESULTS

### Behavioral results

Instructed and Free-Chosen trials were mixed. To determine whether the gestures selected in successive Free-Chosen trials were completely random, we calculated the cumulative number of trials for Power and Hook gestures, respectively. As shown in Figure 3, the curves were very close to the diagonal line for both monkeys, indicating that the two types of trials were conducted equally along the time course. However, this does not exclude the situation that the monkey chose a stereotyped pattern (e.g., continuous Power grip followed by Hook grip). To quantify the randomness of the grasp patterns, we then calculated Shannon’s Entropy index (see methods). If the entropy index was too low, it meant that the monkey chose one of the gestures more often than the other, or the monkey chose the gesture following a stereotyped transition pattern (e.g. Power, Hook, Power, Hook, Power, Hook, etc.). On the contrary, if the entropy index was close to the maximum, it meant that the gesture was chosen irregularly, the outcome of this trial was independent of previous outcomes. In this study, we analyze the 1 to 4 historial trials affect current gesture chose. Firstly, we use 1 historial trial. The number of transition matrix’s states was n=4 ((2×2 matrix, trial l-1×trial l)), the maximum Shannon’s Entropy of 1 historial trial was H_max_=2. Therefore, H_max_=3,4,5 when use 2 to 4 historial trials, and so on. The results of Shannon’s Entropy index were shown in Table 1. This result showed that historial trials did not have much impact on the current choice, the Shannon’s Entropy index were close to the maximum entropy level. These results indicated that monkeys selected Power and Hook gestures in Free-Chosen trials randomly and equally.

**Table 1.** Shannon’s Entropy index of Monkey B06 and Monkey B09

| History trials | 1 | 2 | 3 | 4 |
| --- | --- | --- | --- | --- |
| $H_{\max}$ | 2 | 3 | 4 | 5 |
| Monkey B06 | $1.96 \pm 0.02$ | $2.93 \pm 0.05$ | $3.85 \pm 0.05$ | $4.72 \pm 0.03$ |
| Monkey B09 | $1.97 \pm 0.02$ | $2.89 \pm 0.05$ | $3.81 \pm 0.05$ | $4.69 \pm 0.07$ |

**Figure 3.**
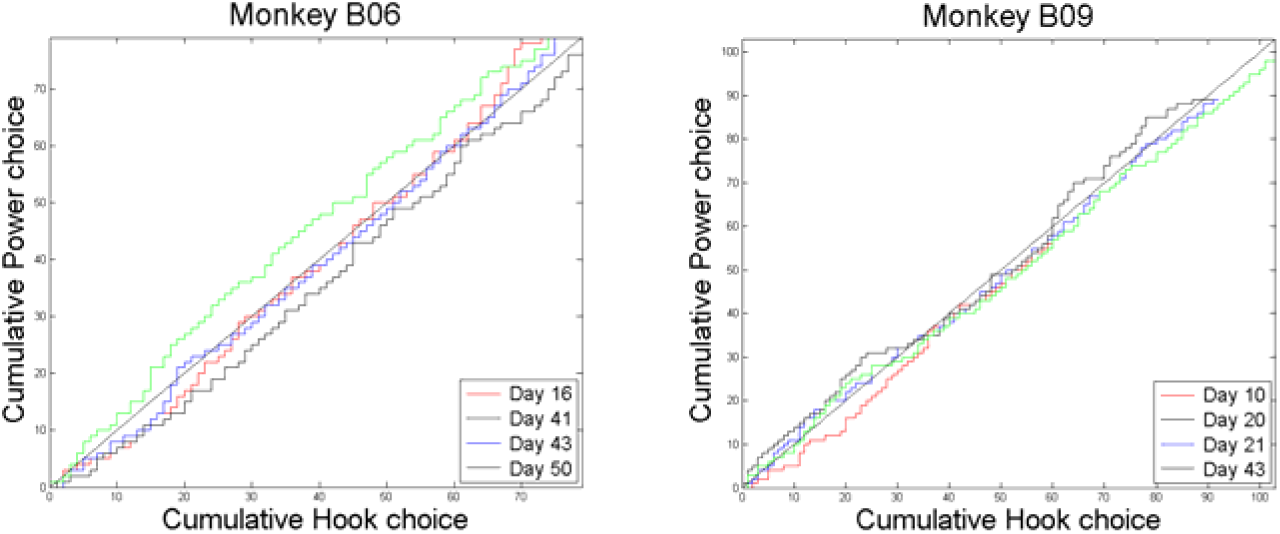
Behavioral choice data from 4 day’s session for each monkey plotting the cumulative number of trials in which the monkeys chose Power and Hook.

### Single-Unit Activity

Figure 4 showed four example units when choosing power and hook gestures in Free-Chosen condition (left) and Instructed condition (right). In general, the firing pattern of these units in Instructed condition looks roughly the same as in Free-Chosen condition. After that, let’s look at these units in detail. The discharge of unit 196 from Monkey B06 (Fig. 4A) and unit 46 from Monkey B09 (Fig. 4B) kept stable before Gesture Cue on, and there was no significant difference between Power and Hook gestures during the baseline period. Because Free-Chosen trials were randomly interleaved with Instructed trials, the condition remained unknown before Gesture Cue on and the monkey should not make any decision in this period. Then the Gesture Cue appeared, neuronal activity diverged after the gestures were chosen. Unit 196 showed selective characteristics of a different choice in Plan period (nearly 700 ms after Gesture Cue on). The firing rate was maintained stable when the monkey chose Power gesture but continued to decrease when the monkey decided to choose Hook gesture (one-way ANOVA, p<0.001). This significant difference was maintained until the movement was executed. Unit 46 illustrated another firing pattern; it showed selectivity immediately after Gesture Cue on. Neuron discharge showed obvious rise when choosing Hook gesture and was significantly higher than Power gesture (one-way ANOVA, p<0.001). Both units showed difference during gesture planning and gesture execution. Unit 182 from Monkey B06 (Figure 4C) only showed significant difference in Plan period (one-way ANOVA, p<0.001), and the firing rate dropped to zero sharply before the movement execution. There were also some purely motor units. Unit 88 (Figure 4D) was a typical unit only showing selectivity during the gesture execution period (one-way ANOVA, p<0.001). In the pre-movement period, this unit was almost inactive. Then we calculated the Pearson correlation coefficient of the average firing rate of these 4 units under both conditions, with r values of 0.98, 0.92, 0.86, and 0.88. It can clear see that the results were all close to 1, The firing pattern of these units is similar under both conditions. It was worth noting that some of these units showed selectivity earlier in Instructed condition (Figure 4A and C).

**Figure 4.**
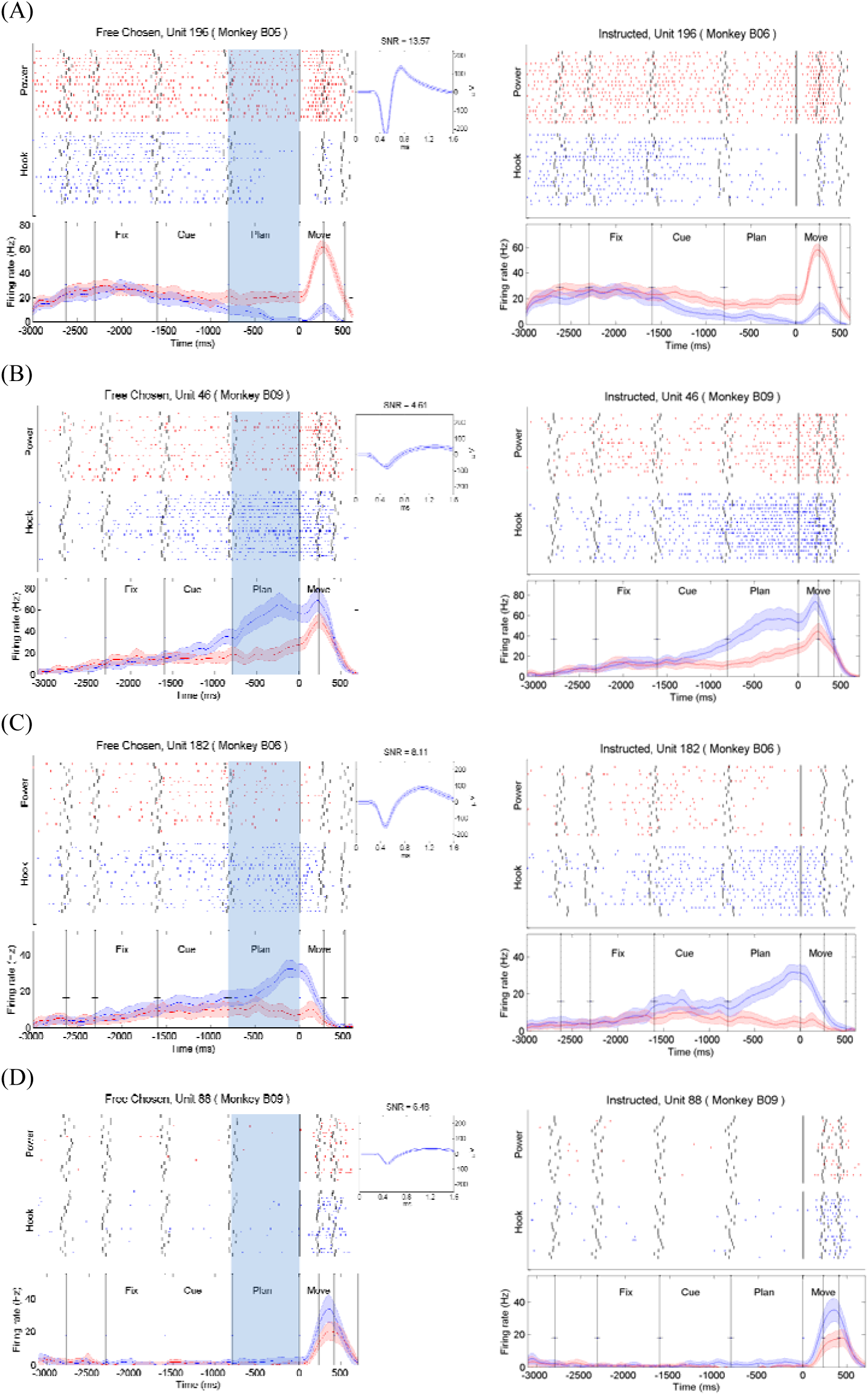
Examples of PMd gesture planning units according to gesture in Free-Chosen condition (Left) and Instructed condition (Right). (A) (B) (C) and (D), examples of PMd gesture planning units. Spikes were aligned on the timing of the Go Cue signal (at 0s). Red (blue) points in raster displays were spike firing time in trials which chose power (hook) gestures. The black vertical ticks in raster displays were behavioral events, from left to right: Button On, Hold-signal On, Gesture Cue On, Gesture Cue Off, Go-signal On, Button Off and Object Sensor On. The seven trial periods are divided by these events. The peristimulus time histograms (PSTH) were smoothed using a Gaussian kernel (SD = 50 ms), and its thickness represents the standard error calculated with the bootstrap method. The waveform and signal-to-noise ratio(SNR) of this unit was shown on the middle.

The four units in Figure 4 were typical in the population of Monkey B06 and Monkey B09. On the basis of unit response properties, the units were subdivided into four main categories: *purely execute* units, *purely plan* units, *plan and execute* units, and *no selective* units. The *purely execute* units only showed selectivity during Move period, such as the case in Figure 4D. The *purely plan* units only showed selective during Fix and Cue period, and not respond during the Move period, such as Figure 4C. The *plan and execute* units showed selective during both Plan and Move periods, such as Figure 4A and 4B. The *no selective* units didn’t show selective during all the trial.

We calculated 3 days data of each monkey, and the statistical results of different kind of units were shown in Table 2. The results showed that In Free-Chosen condition, 9.6% units of B06 and 19.3% units of B09 were *purely plan* units while 15.3% units of B06 and 11.7% units of B09 were *plan and execute* units. It means that nearly 24.9% (B06) and 31.0% (B09) units in PMd participated in gesture planning in Free-Chosen trials. And in Instructed condition, the proportion of *purely plan* units (11.2% of B06 and 20.9% of B09) and *plan and execute* units (17.4% of B06 and 12.0% of B09) were a little more than Free-Chosen condition. Furthermore, we find that 21% units of B06 and 26.8% units of B09 showed selectivity in both Instructed and Free-Chosen conditions (Table 2 overlap). Then we calculated the average correlation coefficient of this part of units. The result showed that units’ trial-averaged responses during the delay strongly correlated for Instructed and Free-Chosen conditions (mean r=0.88±0.12, p < 0.001 for all 6 datasets). Furthermore, all these selective units consistently ‘preferred’ the same gesture in Free-Chosen trials as in Instructed trials.

**Table 2.** The proportion of *purely execute* units, *purely plan* units, *plan and execute* units, and *no selective* units in PMd from 3 day’s session of Free-Chosen and Instructed condition. The third line of each day’s data shows the number of units which illustrated selectivity in both Free-Chosen and Instructed conditions.

| MONKEY |  | Cue | Plan | Execute | Percent of units |  |  |  |  |
| --- | --- | --- | --- | --- | --- | --- | --- | --- | --- |
|  |  |  |  |  | Free-Chosen |  | Instructed |  | overlap |
| Monkey B06 | *plan and execute* | + | + | + | 15.3±1.1% | 7.4±0.6% | 17.4±0.9% | 9.3±2.1% | 12.8±0.9% |
| | | + | - | + | | 1.0±0.3% | | 3.0±1.0% | |
| | | - | + | + | | 7.0±2.8% | | 5.2±0.6% | |
| | *purely plan* | + | + | - | 9.6±2.7% | 3.6±0.7% | 11.2±1.9% | 4.3±0.7% | 8.1±2.2% |
| | | + | - | - | | 2.1±1.4% | | 2.5±0.9% | |
| | | - | + | - | | 3.9±2.3% | | 4.5±2.3% | |
| | *purely execute* | - | - | + | 22.4±2.9% | 22.4±2.9% | 20.0±3.1% | 20.0±3.1% | 18.4±2.6% |
| | *no selective* | - | - | - | 52.7±3.3% | 52.7±3.3% | 51.4±4.7% | 51.4±4.7% | 47.4±3.7% |
| Monkey B09 | *plan and execute* | + | + | + | 11.7±1.6% | 4.7±0.5% | 12.0±2.1% | 5.5±2.6% | 9.6±2.3% |
| | | + | - | + | | 0.4±0.5% | | 1.1±1.6% | |
| | | - | + | + | | 6.6±2.1% | | 5.4±2.0% | |
| | *purely plan* | + | + | - | 19.3±1.1% | 6.6±2.1% | 20.9±2.0% | 7.3±2.6% | 17.2±2.7% |
| | | + | - | - | | 2.2±0.3% | | 3.4±2.1% | |
| | | - | + | - | | 10.6±0.5% | | 10.2±1.1% | |
| | *purely execute* | - | - | + | 4.0±0.5% | 4.0±0.5% | 4.7±1.6% | 4.7±1.6% | 4.0±0.5% |
| | *no selective* | - | - | - | 65.0±5.2% | 65.0±5.2% | 62.3±8.8% | 62.3±8.8% | 59.2±4.9% |

After that, we divided the *purely plan* units and *plan and execute* units into three categories according to the stages of the selectivity that began to appear. The results demonstrated that whether in Free-Chosen conditions or Instructed conditions, the number of units that showed selectivity in Cue period and do not showed selectivity in the Plan period was very few (second row of *purely plan* units and *plan and execute* units in Table 2). In contrast, the number of units that showed selectivity just in Plan period or in Cue and Plan period was more (first and third row of *purely plan* units and *plan and execute* units in Table 2). We observed that most of the units that showed selectivity in the Cue and Plan period tend to maintain selectivity to go-signal.

Figure 5 showed the proportion of gesture planning related units from Monkey B06 and Monkey B09 at different stages of the experiment. Overall, the number of units involved in the gesture planning gradually increased with time (Figure 5A). Figure 5B showed the proportion of selective units in different time in Instructed condition. In this condition, the number of gesture planning units increased sharply and reached to plateau at 400ms after Gesture Cue on. Then the proportion of selective units remained stable unit gesture execution. In addition, there were roughly equal numbers of units which were excited between choosing Power (below the diagonal line) and Hook (above the diagonal line) gestures (number of Power excited units/ number of Hook excited units =1.08±0.21).

**Figure 5.**
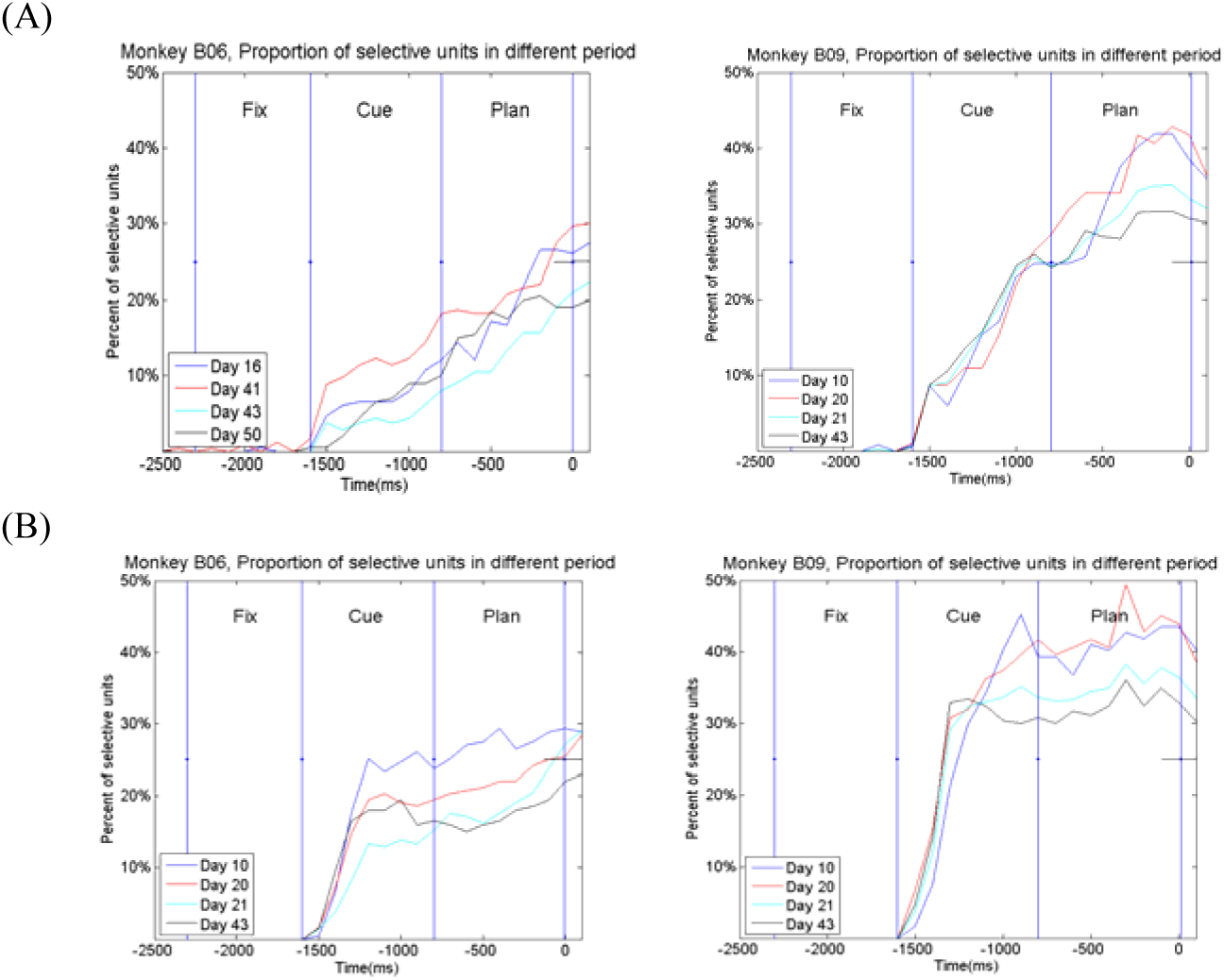
Proportion of selective units in different period. (A) show the proportion of selective units in different time in Free-Chosen condition. (B) show the proportion of selective units in different time in Instructed condition

### Population Analyses

To quantitatively examine the prediction performance of gesture choosing during planning period, we trained an SVM model. The neuronal discharge from 2.7s before Go Signal to 0.6s after Go Signal was employed in the analysis. The result was shown in Figure 6.

**Figure 6.**
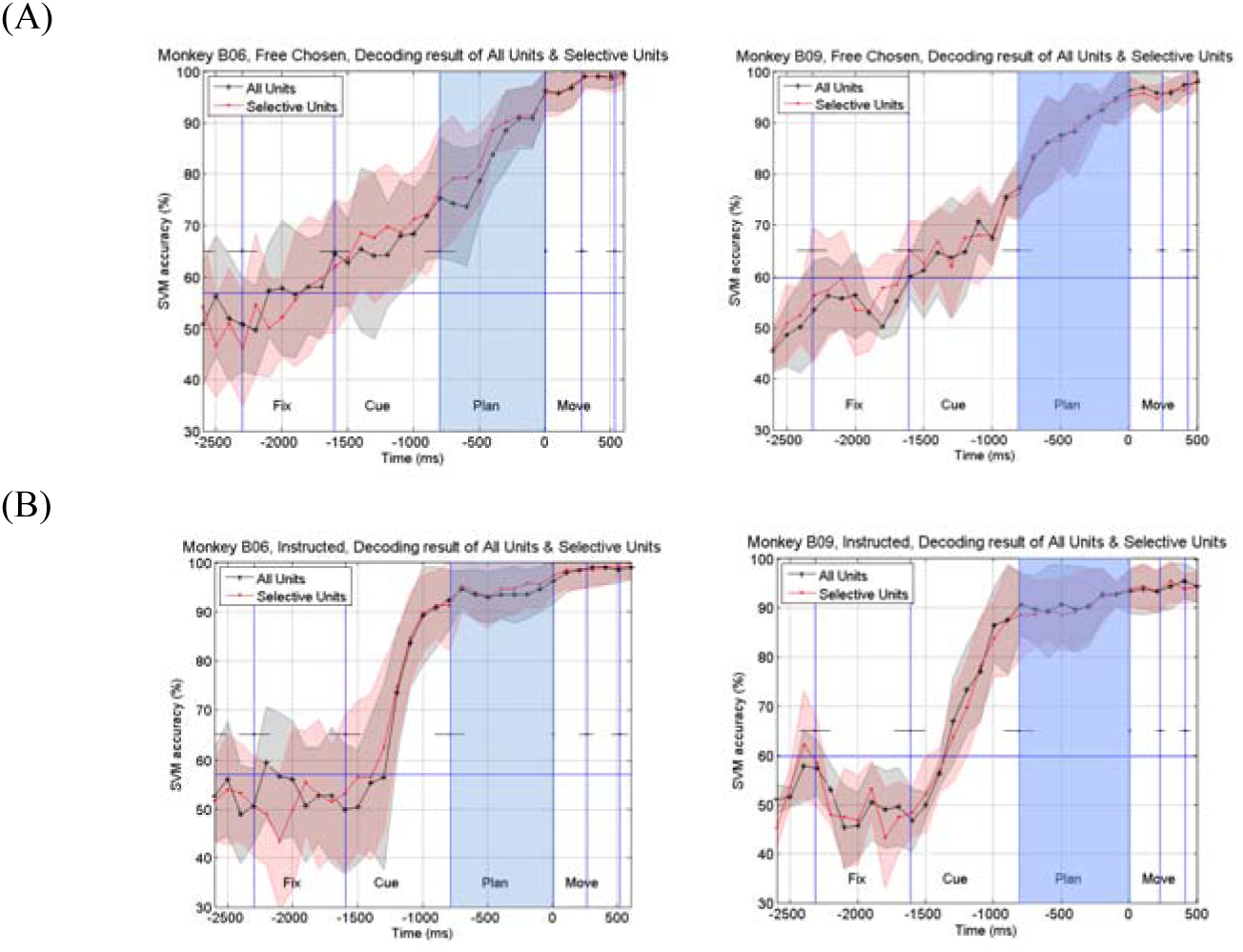
Decoding result analysis. Performance of support vector machine (SVM) decoder using a sliding time window approach. Prediction performance as a function of all units (black line with SEM shadow) and the units tuning to the grip in planning periods (red line with SEM shadow). We use period of 300ms before the time point to classify the grasp gesture. The plots have been aligned at Go Signal as indicated by the vertical lines at 0s point. The blue horizontal line represented the 95% confidence interval of classification from random (57%). The light blue area marks from grip cue onset to Go Signal onset. (A) Decoding result in Free-Chosen condition. (B) Decoding result in Instructed condition.

Prediction performance was shown as two curves: all units (black line) and the units tuning to gestures in planning periods (red line). We used a period of 300ms to classify the grasping gesture, and a 10-folds cross-validation was used. The predicted curves illustrated a monotonic increase in both monkeys. Before the Gesture Cue onset, the accuracy of prediction stuck was at chance level, as the monkey did not make early decision at this period. In Free-Chosen condition (Figure 6A), after Gesture Cue turned on, the accuracy of prediction started to creep up over time. At the end of Plan period, the predicted result reached to 95%. In Instructed condition (Figure 6B), the predicted result raised sharply from 300ms after Gesture Cue on, and at 800ms after Gesture Cue on, the predicted accuracy reached to nearly 90%, then remained stable until the end of Plan period. These results suggesting that we could predict the gesture chosen by the monkey before the movement was executed.

From the previous single neuron analysis and neural cluster analysis, we can see that the units firing patterns were similar under Free-Chosen and Instructed conditions, but the tuning time and cluster coding method may be different. Therefore, we refer to the work of Kaufman et al. and calculated the situation of monkeys changing his mind in some trials (Kaufman et al., 2015). The result was shown in Figure 7. The result of the Free-Chosen condition was shown in left, and the result of the Instructed condition was shown in right. The Y-axis is 1 indicated that the firing pattern of the neural cluster coincided with the time when the Power gesture is about to be performed, and the Y-axis is −1 to indicate that the firing pattern of the neural cluster coincided with the execution of the Hook gesture at this time. The Y-axis at 0 indicates that the firing pattern of the neural cluster is rather chaotic, and the gesture cannot be distinguished. Figure 7 illustrated that a large proportion of trials showed a significant waver between 1 and −1 in Cue and Plan in Free-Chosen condition, they switched between the two gestures until they approached go-signal. In Instructed condition, when Gesture Cue appeared, most of the trials quickly made a choice and kept it until the go-signal appeared, only a few trials showed hesitation. This result is consistent with the previous decoding results.

**Figure 7.**
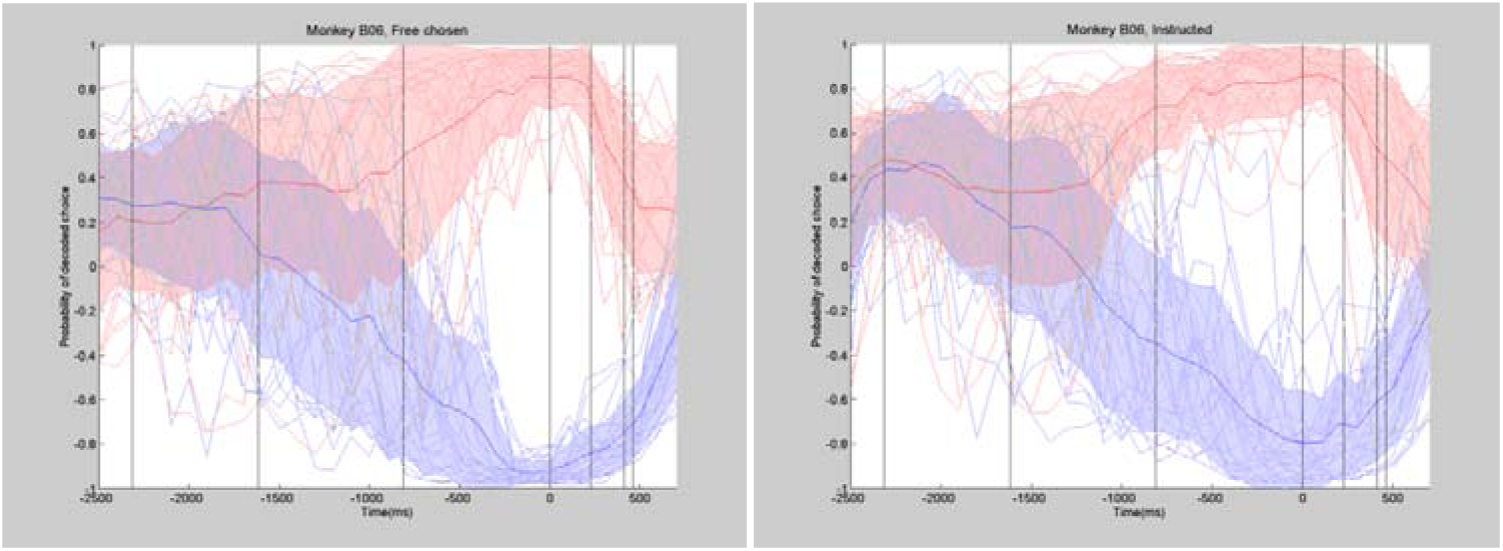
Decoded choice plots for Free-Chosen (left) and Instructed (right) trials, generated by leave-one-out cross-validation. X axis represents time and y axis shows the percentage of fraction correct classification. The light red lines indicate a single trial which chose Power gesture, and light red lines chose Hook gesture. The dark red/blue line with shadow shows the average probability of all trials which chose Power/Hook gesture in Free-Chosen (left) and Instructed (right) conditions.

To visualize the population firing pattern and possible regularity in the data, we used Laplacian Eigenmaps to conduct dimensionality reduction because it encourages the neighbors in the original space to be neighbors in the lower dimensional space. The dimensionality reduction trajectories of each trial in Figure 8 provided clear convergence and divergence of the ensemble neural activity patterns through the time course of planning and execution period. This result illustrated that the firing patterns under each choice were initially similar before gesture cue on. After the gesture cue on, the firing pattern expressed obvious different features and divided into two gesture-related clusters. Finally, the firing patterns of different choices were totally different in execution period. Then we compared the differences in Plan period under the two conditions. In Instructed condition (Figure 8B), the firing pattern divided immediately after the Gesture Cue on. And in Free-Chosen condition (Figure 8A), it showed more confusion in the first half of Plan period. These results meant that the population firing pattern of PMd responded to different gestures in the planning period, and PMd use different strategies to deal with the two conditions.

**Figure 8.**
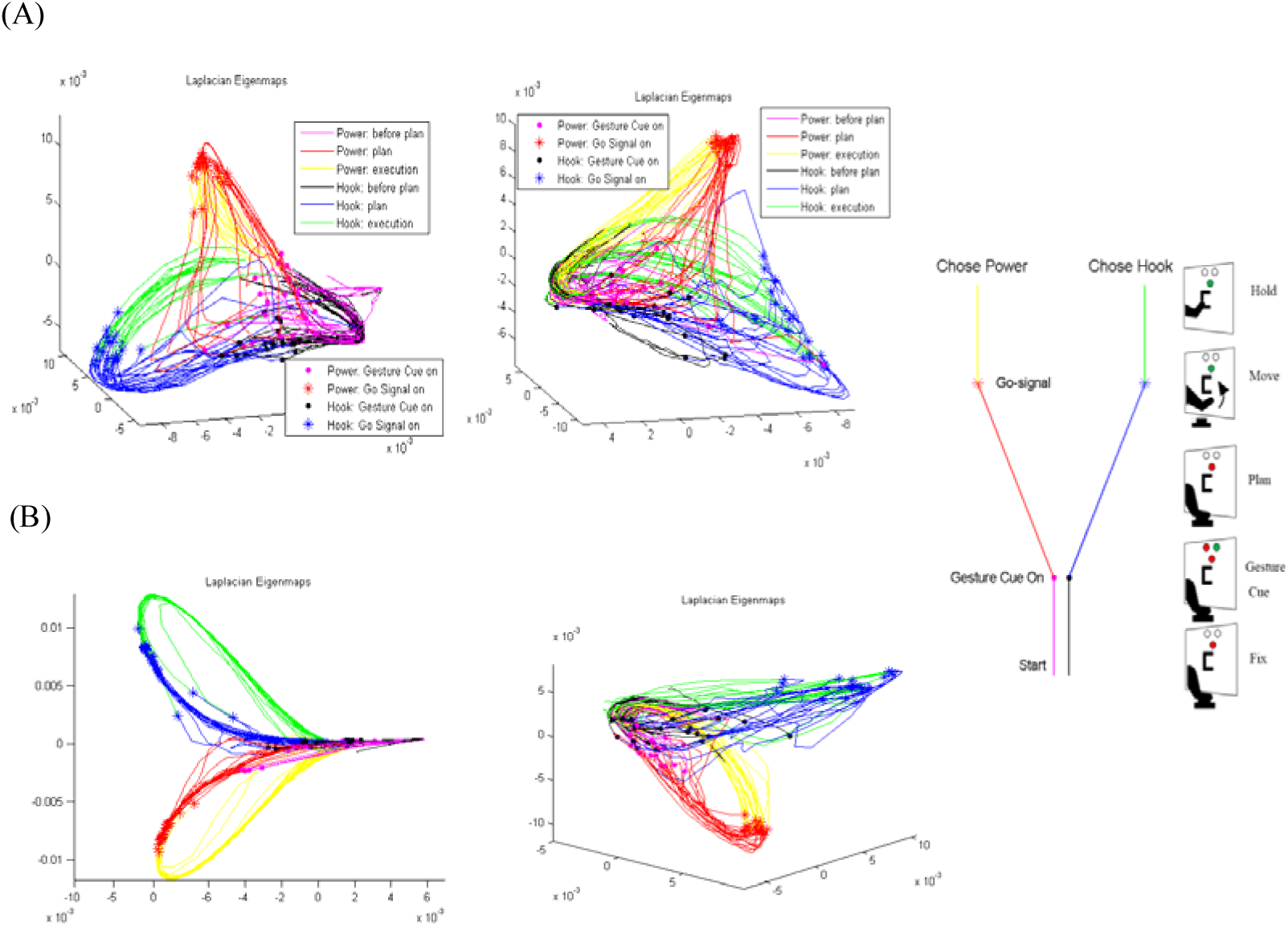
Laplacian Eigenmaps for dimensionality reduction. (A) in Free-Chosen condition and (B) in Instructed condition. We reduce the neural population data of each trial into 3 components, each line represented a single trial. The trial which chose Hook gesture shown in black, blue and green lines, and the trial which chose Power gesture shown in magenta, red and yellow lines. Each color represented different period of this trial, black and magenta illustrated the period from trial on to gesture cue on, blue and red illustrated the period from gesture cue on to go-signal on, green and yellow illustrated the period from go-signal on to trial end. The dot point marked the time of gesture cue on in each trial, and the * point marked the time

To investigate if the selective units for Free-Chosen trials were spatially distributed, we calculated the decoding performance of single unit and plotted on each channel of Blackrock array (Figure 9). We used the data of each single unit to train and test a support vector machine (SVM) model, and if there were multi units in one channel, only the best result was shown. The direction of the wire bundle was marked on the left. Each block in this map represented the location of a channel on Blackrock Array. The four white blocks in each map denoted the channels which were not used for recording. The different color in this map illustrated the maximum decoding result of each single unit on this channel in the planning period. At the same time, we could find that the selective units were distributed evenly in the recording area of PMd in all these data.

**Figure 9.**
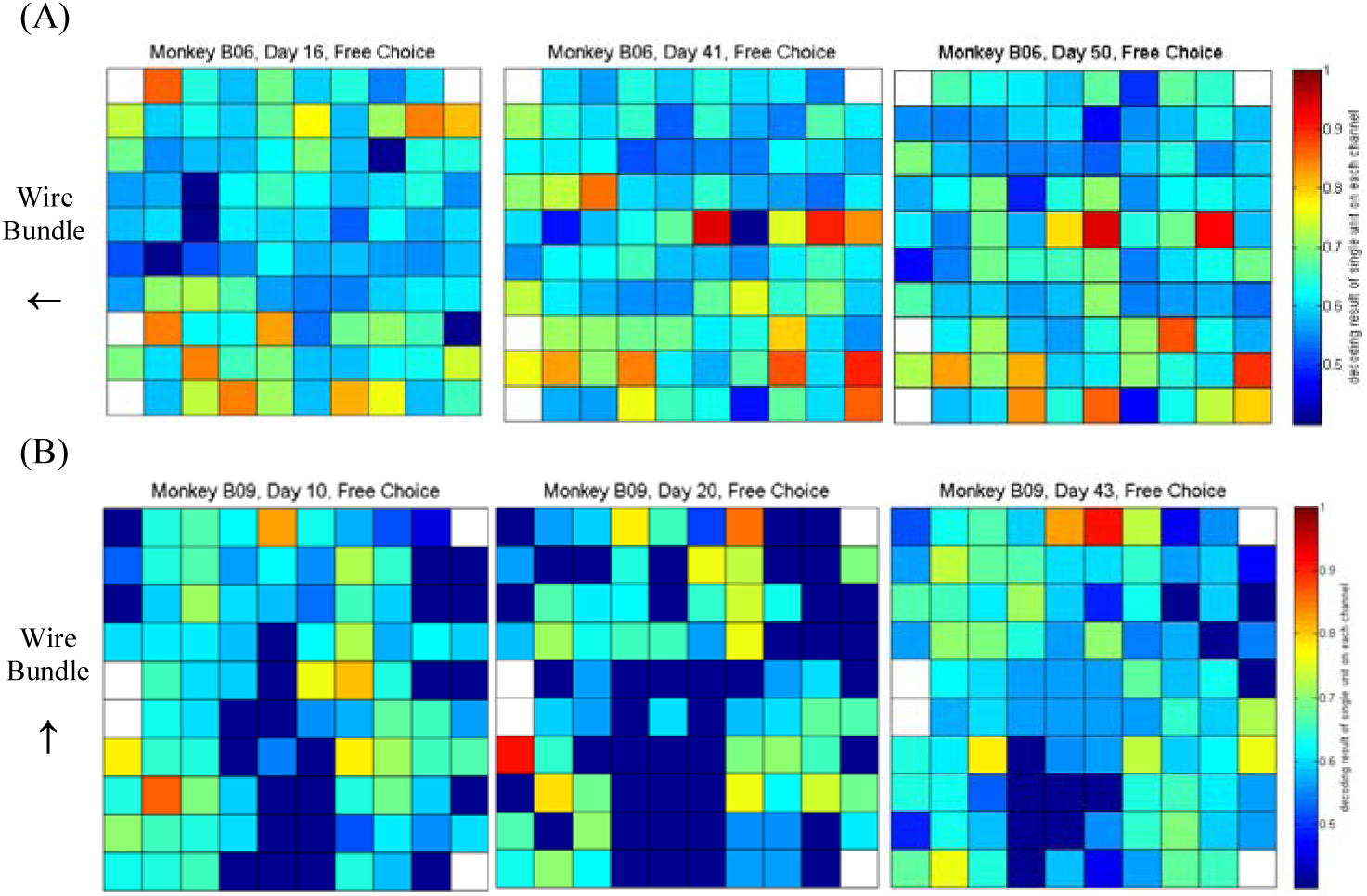
Spatial distribution of selective gesture units mapped by Blackrock Array. (A) and (B) panels are corresponding to Monkey B06 and Monkey B09. We choose 3 different datasets for each monkey, (A) from left to right: the 16th, 41st and 50th day after surgery; (B) from left to right: the 10th, 20th and 43rd day after surgery. Each block in upstairs maps represented the location of a channel on Blackrock Array. The 4 white blocks in each map showed the channels which were not used for recording. The different color in this map illustrate that the maximum decoding performance of single unit on this channel in planning period. The figure downstairs showed the waveform of units on each channel.

## DISCUSSION

### 1. PMd involved in gesture planning

The function of the PMd had been extensively studied for many years. Some works demonstrated that PMd was correlated to parameters of reaching movements which were only involved in proximal forelimb movements (Caminiti et al., 1991; Fu et al., 1993; Kalaska et al., 1997; Wise et al., 1997). These studies had reported that PMd highly participated in modulating reaching. However, Neuro-anatomical evidence that PMv shared reciprocal connections with PMd area (Kurata, 1991; Ghosh et al., 1995; Dancause et al., 2006). Moreover, there were some research reported that reaching and grasping movements planning and execution might share a common brain network (Begliomini et al., 2014; Fattori et al., 2014). In addition, there were some evidences about grasp-related activity recorded in PMd (Raos et al., 2004; Stark et al., 2007; Van et al., 2012; Hao et al., 2013). However, the function of PMd in the grasp planning process is not discussed in depth.

In this paper, we trained the monkeys to grasp same object by freely choosing one of two grips, power grip or hook grip (Cui et al., 2007; Michaels et al., 2015; Dann, 2017). The results showed that 21.0% or 26.8% units in PMd of each monkey showed selectivity for different gestures (one-way ANOVA, p<0.01) before movement executed. This observation provided direct evidence that the PMd participated in gesture planning.

It was worth noting that, the biggest difference between the two gestures was that in Power gesture the thumb was opposite to the other four fingers, and in Hook gesture thumb was close to the other four fingers. Therefore, it can also be considered that these selective units of PMd were sensitive to the activity of the thumb because the thumb was 2 degree-of-freedom and had a greater influence on the gesture (Santello et al., 1998, 2002). In addition, the number of units from PMd excited by Power gestures was approximately equal to the number of units excited by Hook gestures (Figure S1). PMd did not show preference to a specific action.

In addition, to ensure that the selectivity of PMd is indeed related to gestures but arm position, we designed a complementary experiment to verify it. The result showed that the small differences in arm positions between different gestures do not cause the selectivity in PMd (See Supplementary).

### 2. The neuron selectivity in planning period was internal generated not external visual stimuli

Recent years, many studies had focused on the visual guidance of motor behavior function of PMd (Johnson et al., 1996; Wise et al., 1997; Hoshi and Tanji, 2007; Averbeck et al., 2009). These studies concluded that PMd was a key node of network underlying visually guided reaching (Kalaska and Crammond, 1995; Johnson et al., 1996). On the other hand, some studies showed considerable evidence that the function of control sequential movements were internally generated (Kurata and Wise,1988; Mitz et al., 1991; Cisek and Kalaska, 2005; Ohbayashi et al., 2016).

In the classical experiment of turning table, the choice of gesture is related to the information of the specific object such as the size and shape of the object. In our and Scherberger’s experiment (Michaels et al., 2015; Dann, 2017), the monkeys faced the same object, and the hint information was an abstract visual cue. Therefore, the selectivity of PMd is not related to the characteristics of the object.

Was the selectivity of PMd related to visual cue or hand gesture? In our Free-Chosen condition, there was no difference between visual cue, and part of PMd units showed selectivity of different gestures. This kind of selectivity cannot be induced by external visual stimuli. Furthermore, in the Instructed condition, the monkeys face only red LED or only greed LED during the Cue period, and the monkeys face the red and green LED under the Free-Chosen condition. Therefore, if PMd responded to visual stimuli, there should be some neurons in the Cue period that showed selectivity only in Instructed conditions and showed no selectivity in Free-Chosen conditions (Ariani et al., 2015). However, in Table 2 we could hardly observe such units, so we think that these selectivity units of PMd might participate in internally driven gesture planning.

### 3. Internally driven gesture planning more inclined to coding gesture movement preparation

Regarding the function of premotor area in the process of internally planning, some scholars reported that it had the function of decision making. Cisek and Kalaska reported that PMd may eventually eliminate some possible options through information accumulates and make a final decision (Cisek and Kalaska, 2005). Donner et al. showed that during motion detection, motor plans for both “yes” and “no” choices result from continuously accumulating sensory evidence (Donner et al., 2009). However, there were also researchers reported that PMd had the function of linked between decision making and action (Svoboda et al., 2017; Churchland et al., 2006; Roitman et al., 2002). In addition, some scholars demonstrated that the function of PMd was related to the abstract movement (Kaufman, 2015; Ariani et al., 2015). We can clear see that the changes in the number of selective units (Figure 5) and decoding results (Figure 6) showed that PMd quickly chose the final gesture and kept it in motion in Instructed condition. However, monkeys need more time to make final decision. It was clear that monkeys use different selection strategies under Instructed and Free-Chosen conditions. But the final output action is the same (Power or Hook). Therefore, if in these two conditions, the individual and cluster firing patterns of PMd units are closer, the function of PMd is more inclined to gesture movement preparation, otherwise, the function of PMd is more inclined to selection strategies. In our results, we found that most of selective units showed selectivity in both Instructed and Free-Chosen conditions, and the correlation coefficient result showed that units’ trial-averaged responses during the delay strongly correlated for Instructed and Free-Chosen conditions (mean r=0.88±0.12, p < 0.001 for all 6 datasets). Furthermore, all these selective units consistently ‘preferred’ the same gesture in Free-Chosen trials as in Instructed trials. Therefore, we consider that the function of this part of PMd neurons is more inclined to gesture movement preparation.

From Figure 7 and Figure 8, some trials had oscillated during the Cue and Plan period. This result indicated that the monkeys may hesitate during the planning period (Kaufman, 2015). On the one hand, this result showed why the decoding performance was slowly increasing in Free-Chosen condition (Figure 6). On the other hand, the SVM using was trained by using only Instructed trials with delay durations of at least 300 ms. This part of neural activity should be highly related to the gesture movement preparation. In the Free-Chosen condition, the neural decoding results fluctuated greatly between 1 and −1, rather than slightly swinging around 0. This result illustrated that the selective neuron firing mode of PMd is highly correlated with the gestures movement preparation from another side. This result was coincided with fMRI result of human being (Ariani et al., 2015).

From our results, we consider that PMd did not participate in action decision, but only received the decision from upstream brain area. Neuro-anatomical evidence that PMd received projections from frontal PFC, SMAs, PMv and MIP, which upstream brain region driven the decision-making strategies remains to be further studied.

## CONCLUSION

This article focuses on the function of PMd in the grasping planning. The most important observation of this study was that nearly 21.0% and 26.8% units in PMd of two monkeys displayed gesture selectivity during gesture planning in both Instructed or Free-Chosen conditions. These units exhibit selectivity for different gestures when facing the identical visual stimuli (freely choosing condition). At the same time, similar activity patterns are displayed for the same gesture when faced with different selection strategies (freely choosing condition vs. instructing condition). These results show that some neurons of PMd are mainly involved in the hand shape preparation and have no obvious relationship with external visual stimuli and selection strategies.

