## Supplementary for "Dorsal Premotor Cortex Involved in Hand Gesture Preparation in Macaques"

In order to analyze whether the selectivity of PMd is related to different gestures or different positions of the arms, we analyzed the wrist orientation of Power gestures and Hook gestures first. We used motion capture to record the wrist orientation of the two gestures and align the different trials by the center point. The results were shown in Figure S1. It can be seen that the wrist orientation of the two gestures is basically the same. After that, we performed t-test on the two direction vectors of Power and Hook gestures, and the results showed that there was no significant difference between the two groups of data (p> 0.05).

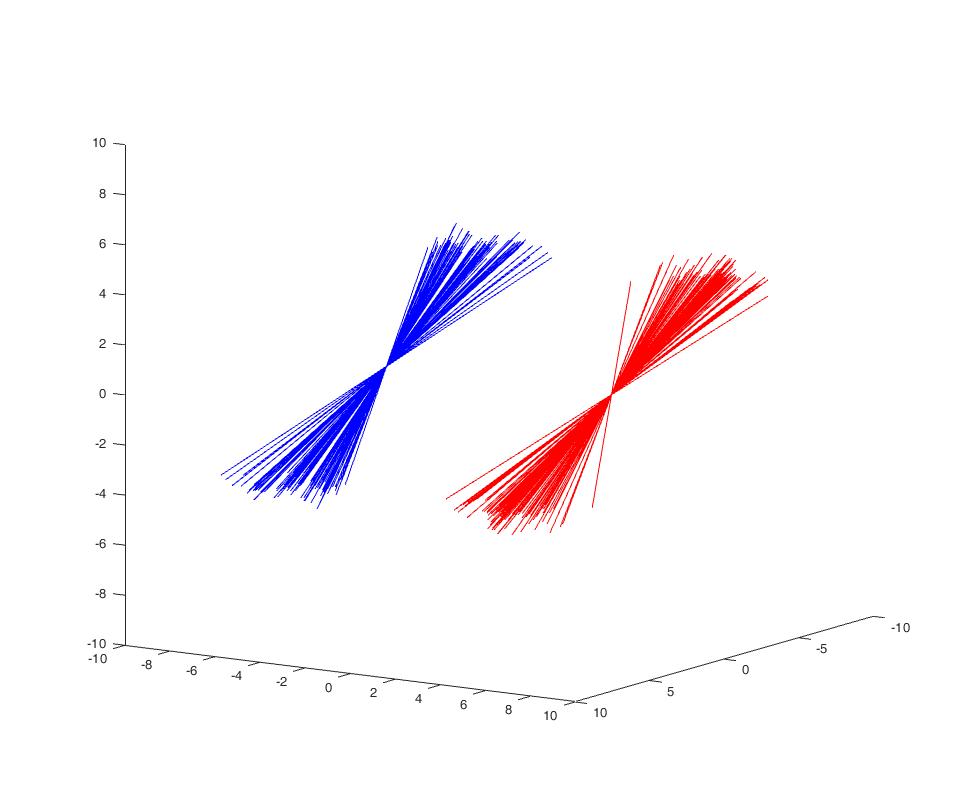

Figure S1. Wrist orientation statistics of Power and Hook gestures. We attached two markers in the vertical direction of the monkey's wrist, and observed the orientation of the wrist through its connection line. The red line indicates the wrist orientation of Power gesture, and the blue line indicates the wrist orientation of Hook gesture. The different trials are aligned by the center point of each line.

Then we analyzing the position of the wrist, and found that the distance of the wrist between the two gestures was about 0.7cm. In order to observe whether the difference of wrist position would lead to the selectivity of PMd, we designed the following experiment: At the end of each trial, the object platform will randomly move to one of the following seven coordinate points, namely Middle (0,0,0), Front (0,1,0), Back (0,- 1,0), Left (-1,0,0), Right (1,0,0), Up (0,0,1) and Down (0,0, -1), the unit of the axis is centimeter. Then, the Support Vector Machine decoding analysis was performed on the x-axis (Left, Middle, Right), y-axis (Front, Middle, and Back) and z-axis (Up, Middle, and Down). The results were shown in Figure S2. It can be seen that the tiny change of 1cm in the x-axis, y-axis or z-axis direction will not cause the selectivity of PMd, but the selectivity between different gestures is obvious, so it can be explained that the selectivity of PMd not caused by the small fluctuations in the arm position but related to the difference of hand gesture.

(A) (B)

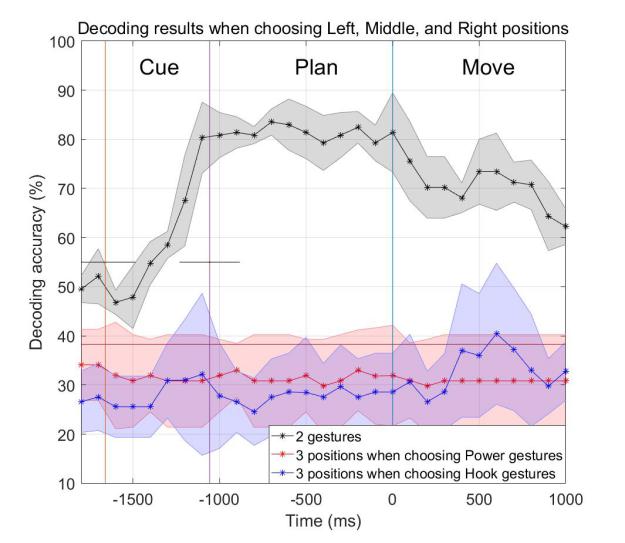

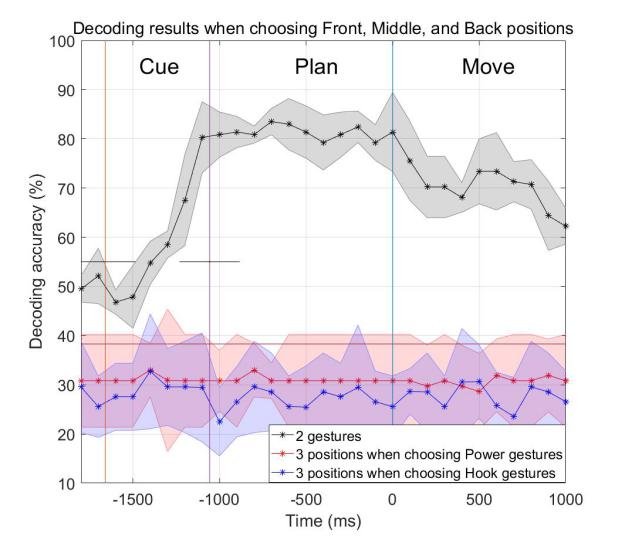

(C)

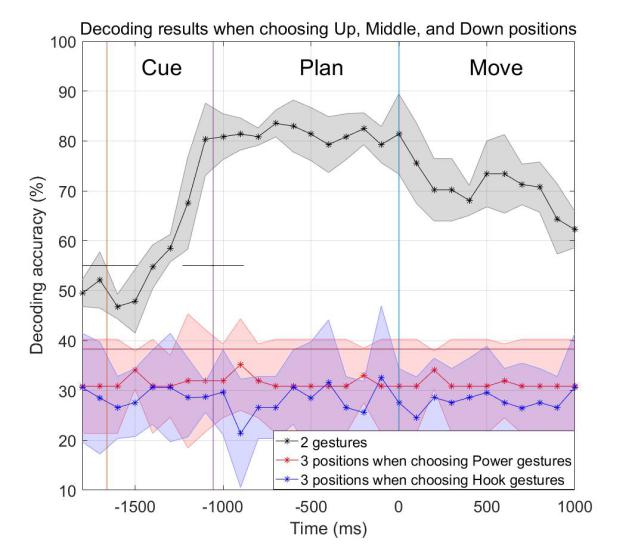

Figure S2. Decoding results of different wrist positions and hand gestures. (A) Left, Middle and Right (B) Front, Middle and Back (C) Up, Middle and Down
